# Changes in biodiversity impact atmospheric chemistry and climate through plant volatiles and particles

**DOI:** 10.1101/2023.01.28.526055

**Authors:** Anvar Sanaei, Hartmut Herrmann, Loreen Alshaabi, Jan Beck, Olga Ferlian, Khanneh Wadinga Fomba, Sylvia Haferkorn, Manuela van Pinxteren, Johannes Quaas, Julius Quosh, René Rabe, Christian Wirth, Nico Eisenhauer, Alexandra Weigelt

**Affiliations:** Institute of Biology, Leipzig University, 04103 Leipzig, Germany; Leibniz Institute for Tropospheric Research (TROPOS), Atmospheric Chemistry Department (ACD), 04318 Leipzig, Germany; German Centre for Integrative Biodiversity Research (iDiv) Halle-Jena-Leipzig, 04103 Leipzig, Germany; Institute for Meteorology, Leipzig University, 04103 Leipzig, Germany

**Keywords:** Biodiversity, biogenic aerosol, BSOA, biosphere-atmosphere interactions, biotic and abiotic stress, BVOCs, climate change, volatile organic compounds

## Abstract

Climate extremes in tandem with biodiversity change affect emissions of biogenic volatile organic compounds (BVOCs) from plants and, as a result, the formation of biogenic secondary organic aerosols (BSOA). The resulting BSOA can have a wide variety of impacts, such as on Earth’s radiative balance or cloud- and precipitation formation. However, at present, it is unclear how changing biodiversity will lead to changes in BVOC emissions, BSOA formation and their corresponding effects. We present a conceptual framework of the relationships between biodiversity and BVOC emissions based on our current mechanistic understanding and combining knowledge from the fields of biology and atmospheric chemistry. Parts of this framework are tested in a case study using a tree diversity experiment with adjunct BVOC and BSOA characterisation. The relative differences in tree monocultures and mixtures show that the overall concentration of BVOCs decreases with increasing biodiversity (*p* < 0.01), but results for BSOA compounds are mixed and overall non-significant (*p* = 0.40). We suggest future studies should follow a multidisciplinary approach where the fields of biology, atmospheric chemistry and climate research interact.

## Introduction

Biosphere and atmosphere are tightly interconnected ^1,2^, making the understanding of their interactions critically important as human life depends on them ^3,4^. On the one hand, a suite of atmospheric drivers can affect the biosphere in different ways ^2^, including detrimental impacts of rising temperatures on ecosystem integrity and biodiversity ^5^. On the other hand, the biosphere can exert feedbacks to the atmosphere, e.g. by releasing various biogenic volatile organic compounds (BVOCs) through plants ^6,7^ or soil ^8^. Following their emission, the atmospheric oxidation of plant-emitted BVOCs leads to gas-phase products, which can either condense on already existing particles or form new ones ^9^, resulting in biogenic secondary organic aerosols (BSOA) ^10^, which are important particles for the radiative balance of the Earth ^11^. Climate extremes as well as biodiversity change affect emissions of BVOCs from plants ^8,12^ and, consequently, BSOA formation ^13^. However, studies using concerted measurements of BVOCs and BSOA along diversity gradients of emitting plant species are largely missing ^14^. This lack of knowledge is particularly worrying given that human activities have traditionally favoured monoculture plantations ^15^, and scientific guidance is urgently needed for major reforestation efforts, especially in the light of current global climate change ^16^.

Tree species release a large variety of hydrocarbons, especially isoprene for deciduous trees and monoterpenes for coniferous trees. However, the composition and magnitude of emitted BVOCs are highly species-specific ^17,18^. These BVOCs are produced primarily via the leaf surface as a result of metabolic processes and as a reaction to abiotic and biotic stress ^19,20^. Plant-released BVOCs determine important biotic interactions such as intra- and interspecific communication ^21^ as well as herbivore and pathogen defences ^22^. BVOCs can also protect plants against abiotic stress such as heat, drought, and high radiation ^20,23^.

In the forest, individual trees are not isolated but compete with their con- and heterospecific neighbours for resources, such as light, water, and nutrients ^24^. Resource availability directly affects eco-physiological processes such as photosynthetic rates ^25–27^ and thus allocation to growth and potential interaction strength. In monocultures or low diverse forest stands, neighbouring trees share largely identical ecological niches and thus strongly compete for available resources. In contrast, mixed forests enable resource partitioning, e.g. via differences in tree architecture ^28^ or rooting depth ^29,30^, often leading to higher stand productivity ^31^ and leaf area index ^32^. Likewise, forests with diverse leaf chemical traits can enhance soil nutrient availability through diverse plant inputs of leaf and root litter and through soil microbial communities and activities ^33,34^, thereby increasing forest stand productivity. As forest biomass production and BVOC emissions are highly correlated ^35,36^, diverse forests should thus emit higher amounts of BVOC (Figure 1, Hypothesis 1).

**Figure 1.**
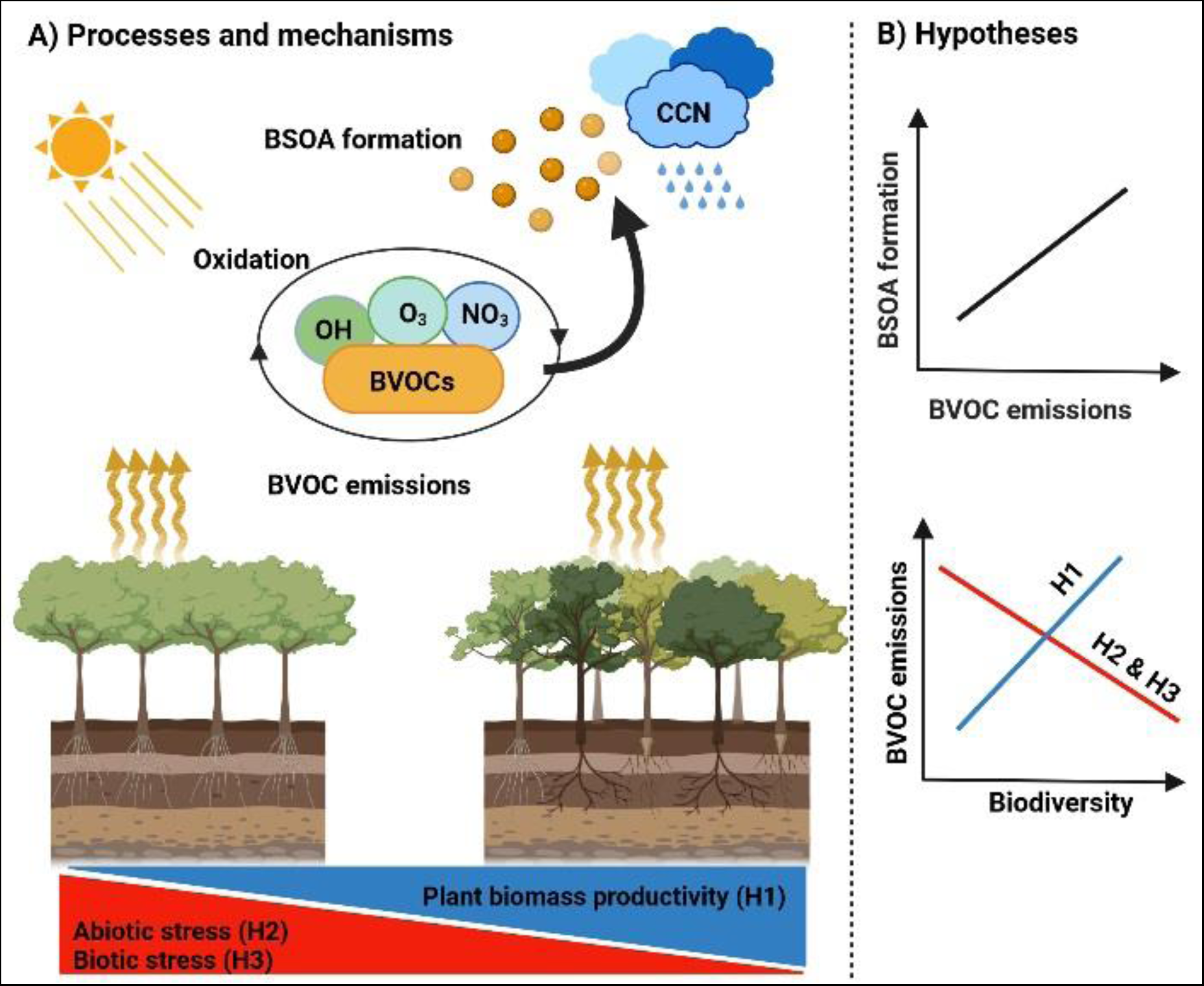
The figure outlines (A) the processes and mechanisms, and (B) the general hypotheses related to biogenic volatile organic compound (BVOC) emissions across biodiversity gradients (monoculture vs. mixtures), biogenic secondary organic aerosol (BSOA), and cloud condensation nuclei (CCN) formation. The hypothesized positive and negative relationships between BVOC emissions and biodiversity are represented by blue and red lines, respectively. The hypothesized positive relationship between BVOC emissions and BSOA formation is indicated in black.

Yet, there are at least two counteracting mechanisms which should lead to decreased BVOC emissions with increasing tree diversity. First, more diverse plant communities increase facilitation and thereby might mitigate abiotic stress. It has been shown that functionally diverse plant communities can increase water use efficiencies through variation in root architecture as a result of spatiotemporal complementary resource use ^29^ and microclimate amelioration, particularly in dry seasons ^37^. In addition, complexity in the vertical and horizontal stratification of diverse stands can relieve thermal stresses ^38,39^ and alleviate the risks of drought stress ^40^. Indeed, decreasing heat and drought stress in mixed communities is likely to result in lower leaf temperature, which could result in lower BVOC emissions ^41,42^. Given that abiotic stress, but particularly heat and drought are known to be important drivers of BVOC emissions, we hypothesize a decrease in BVOC concentrations in mixtures where abiotic facilitation mediates abiotic stress reduction (Figure 1, Hypothesis 2).

Second, more diverse plant communities show decreased per capita herbivory damage and pathogen infection ^43,44^. Although multi-plant diets may be beneficial for different herbivores *per se* ^45,46^, plant natural enemies are also more abundant in more diverse plant communities, which could reduce herbivory pressures as well as insect outbreaks (top-down regulation), as supposed by the “enemies hypothesis” ^44,47^. Furthermore, the heterogeneity of plant nutritive traits in diverse stands suppresses herbivore abundance, resulting in declining herbivore performance ^48^, likely due to non-host species reducing the accessibility of host species ^49^. Overall, reduced biotic stress through herbivores and pathogens in mixtures should reduce the amount of BVOC emissions (Figure 1, Hypothesis 3) but might increase the diversity of emitted compounds. However, irrespective of the change in the amount of BVOC emissions with increasing tree diversity – which might increase or decrease depending on the primary mechanism – we would expect that the diversity of emitted compounds increases in mixed forests given that BVOC emissions are known to be highly species-specific ^50^.

Once entering the atmosphere, plant-derived VOCs react quickly with ambient hydroxyl radicals (OH), Ozone (O_3_), and nitrate radicals (NO_3_) resulting in the formation of BVOC oxidation products that are more oxidized than the compounds originally emitted. These compounds, including highly oxygenated molecules (HOMs) or, as more recently called, oxygenated organic molecules (OOMs) ^51,52^, have lower vapor pressures and better water solubility and can hence contribute to BSOA as atmospheric particles ^53^. Changing BSOA fractions in organic particle composition could have different effects. First, it could influence radiative effects of the particles themselves (‘direct effect’) and hence couple to climate. Secondly, such compositional changes could influence the particles’ ability and effectiveness to act as cloud condensation nuclei (CCN), thus influencing cloud formation with all its respective consequences for radiation through aerosol-cloud interactions (‘indirect effects’) that also affect precipitation formation rates ^54–56^. However, the conversion of BVOC to BSOA is strongly determined by the local atmospheric composition and oxidation regime, temperature, the kinetics and mechanism of the emitted BVOCs and their respective reactions in the gas phase including their phase transfer characteristics and their particle-phase chemistry ^57–59^. High BVOC emissions, e.g. due to rising temperature, increase BSOA formation which, to some extent, could trigger a heating by direct particle absorption but it could also lead to a cooling due to the effect of BSOA on cloud formation and reflectivity of clouds ^11,13^. Recent findings suggest that biotic and abiotic stress-induced BVOC emissions even accelerate climate-relevant BSOA formation ^60–62^.

Our mechanistic understanding and existing knowledge suggest that increasing biodiversity could predictably influence the amount of BVOC released. So far, studies have primarily focused on single tree species though, with no studies available at the time using concerted measurements of BVOCs and BSOA in tree stands or along experimental or natural tree diversity gradients (Figure 1). We argue that this is a critical knowledge gap, as BVOC composition has been recognized as a powerful stress indicator and an important feedback mechanism of climate change ^63^. Addressing this knowledge gap is relevant to understand the causal connections between biodiversity and climate change and to overcome these coupled crises as two of the most urgent challenges facing humanity ^64^. To support our claim that biodiversity might play a significant role in this context and to inspire future research, we present the first data from a case study in the MyDiv tree diversity experiment site in Germany ^65^. Here, we simultaneously measured the magnitude and variability of BVOC and BSOA compounds in ten plots differing in tree diversity. Using this case study, we aim to put forward the link between biodiversity and BVOC/ BSOA and provide a first test of principle in support of the conceptual framework. Our results show that the amount of BVOCs tends to decrease with biodiversity in most cases, while mixed results were found for BSOA compounds. Our findings highlight the need for multidisciplinary work at the interface between the biosphere and the atmosphere to better understand the reciprocal effects of biodiversity and climate change ^4^.

## Results and discussion

### BVOC emissions across biodiversity

We quantified nine different BVOCs from the investigated plots in the MyDiv experiment, i.e. α-pinene, camphene, β-pinene, Δ3-carene, p-cymene, limonene, α-terpinene, isophorone, and acetophenone. Our results show that most of the observed BVOC compounds were not significantly different between the individual tree species in monocultures and their corresponding mixtures, with some notable exceptions (Figure 2). In particular, we find some significant differences in the emission of limonene and acetophenone (ANOVA: *p* = 0.01 and *p* < 0.01, respectively; Figure 2A,I) which was higher in monoculture plots, particularly in *Sorbus aucuparia* L. (Figure 2A,I), highlighting the species-specific dependency of BVOC emission rates ^17,18^.

**Figure 2.**
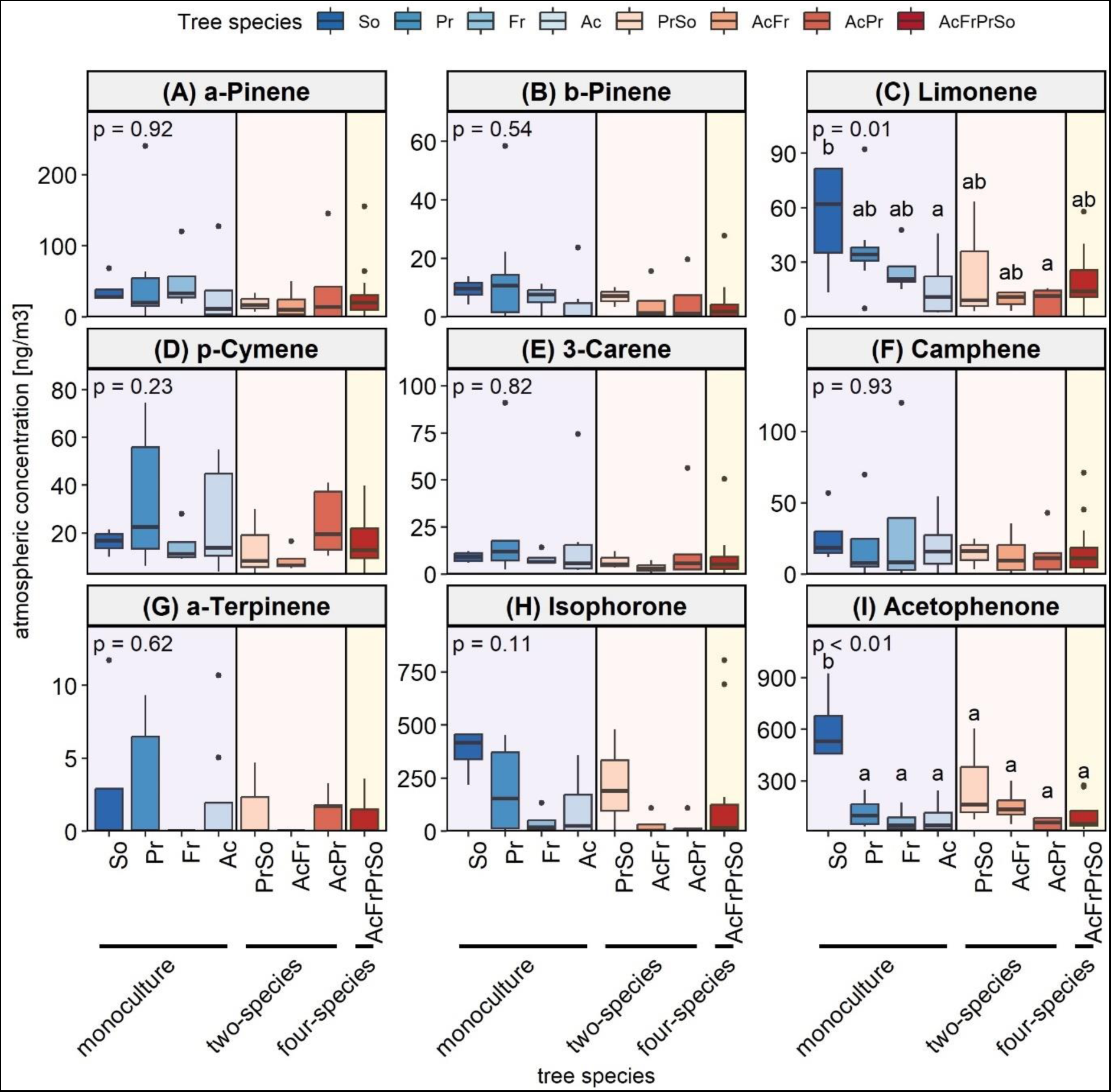
Changes in observed emitted BVOC compounds across individual tree species and mixtures. Boxes indicate interquartile ranges (bottom and top parts of the box), and median lines (horizontal lines within boxes) and whiskers indicate the minimum and maximum of the measurement. *p*-values are shown. Different letters above boxes represent significant differences among tree species (*p* < 0.05; Tukey’s test). Abbreviations: So, *Sorbus aucuparia* L.; Pr, *Prunus avium* (L.) L.; Ac, *Acer pseudoplatanus* L., and Fr, *Fraxinus excelsior* L.

The comparison between the observed and expected amounts of each BVOC compound in mixture plots (two- and four-species mixtures) shows that there were no significant differences; however, we found a few exceptions (Figure 3). In four-species mixtures, some BVOC compounds differed significantly between the observed and expected amounts (Figure 3). As such, the observed amounts of limonene and acetophenone were significantly lower than the expected values (t-test: *p* = 0.04 and *p* = 0.01, respectively; Figure 3C,I). This indicates that four-species mixtures produced lower amounts of limonene and acetophenone than expected.

**Figure 3.**
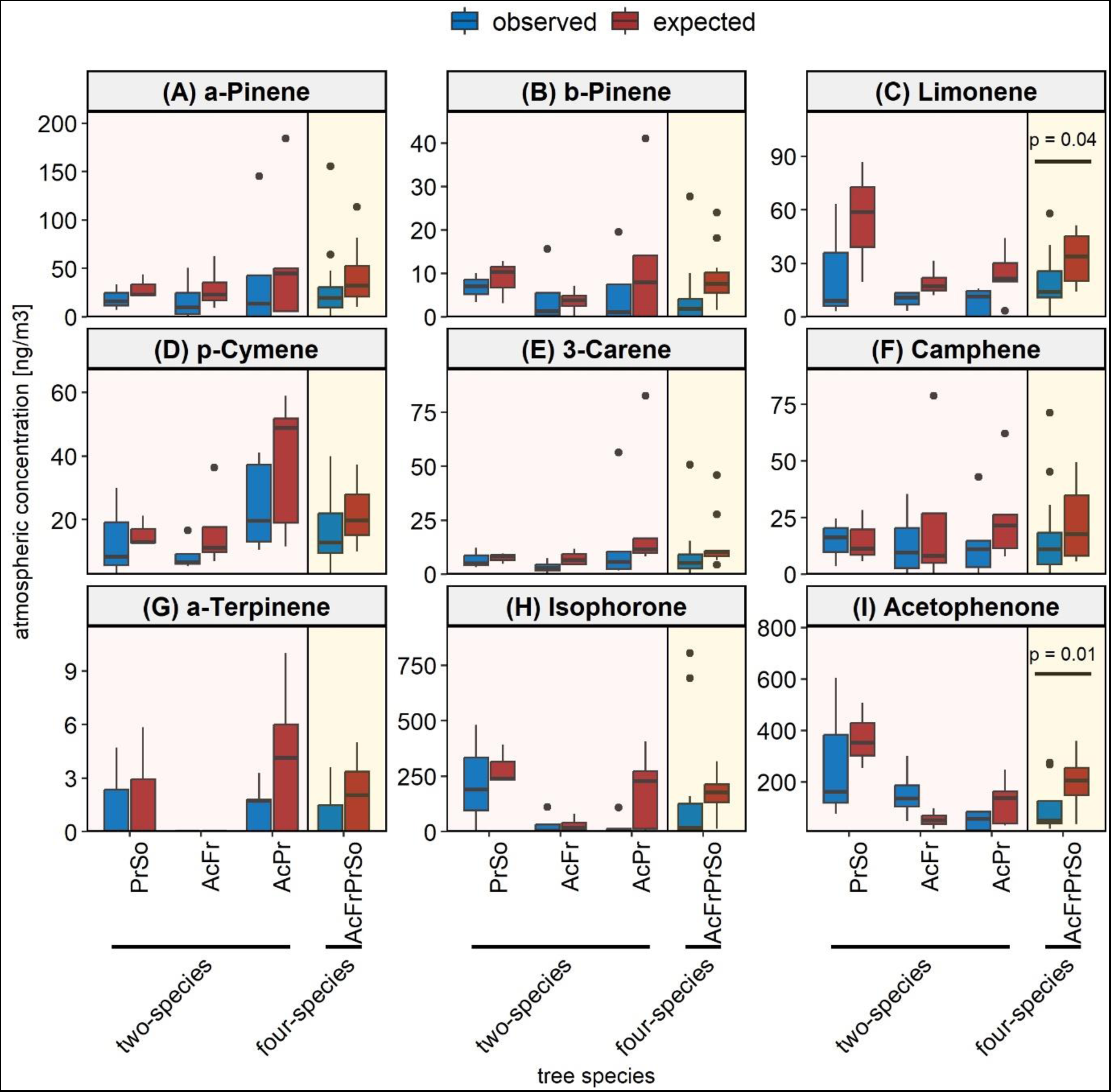
Changes in observed and expected BVOC compounds across tree mixtures (two- and four-species mixtures). Boxes indicate interquartile ranges (bottom and top parts of the box), and median lines (horizontal lines within boxes) and whiskers indicate the minimum and maximum of the measurement. *p*-values are shown (t-test). Abbreviations: So, *Sorbus aucuparia* L.; Pr, *Prunus avium* (L.) L.; Ac, *Acer pseudoplatanus* L., and Fr, *Fraxinus excelsior* L.

The relative differences in monocultures and mixtures based on standardized mean difference analysis across diversity levels show that increasing tree diversity significantly decreased the overall concentration of BVOCs (the overall standardized mean difference: *p* < 0.01, Figure 4). Particularly, the relative differences in the means of observed and expected amounts of limonene were significant (the standardized mean difference: *p* < 0.01, Figure 4), and p-cymene and α-terpinene were marginally significant (the standardized mean difference: *p* = 0.09 and *p* = 0.07, respectively; Figure 4), meaning that mixtures produced significantly less than expected amounts of these compounds compared to monocultures. The relative differences in monoculture and mixtures for two-and four-species mixtures also showed the same decreasing trend of BVOC emissions with increasing tree diversity and were significant (the standardized mean difference, *p* < 0.01, Supplementary Figure 1). As such, the relative differences in the means of observed and expected amounts of limonene were marginally significant in two-species mixtures (the standardized mean difference: *p* = 0.06, Supplementary Figure 1A), while acetophenone and α-terpinene showed strong and marginal significance in four-species mixtures, respectively (the standardized mean difference: *p* = 0.02 and *p* = 0.09, respectively; Supplementary Figure 1B). The overall results indicate that tree diversity significantly reduced BVOC emissions, supporting our second and third hypotheses (H2 and H3).

**Figure 4.**
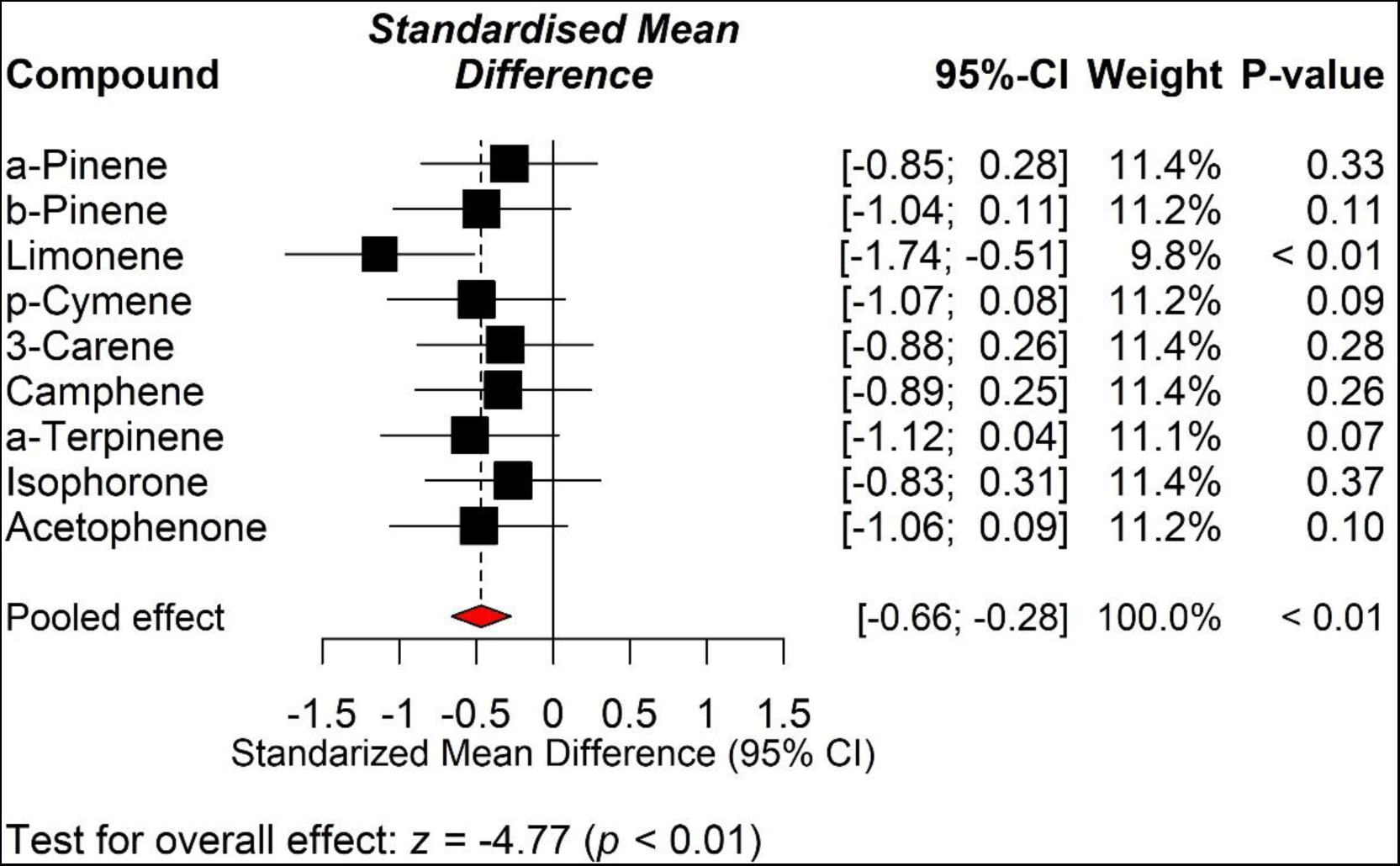
A forest plot summarizing the results of the standardized mean difference between monocultures and mixtures for changes in BVOCs as a result of diversity change. The squares’ sizes and widths represent each compound’s weight, while the diamond (red colour) represents the overall effect estimate of the analysis. Effect sizes were calculated using the standardized mean difference. The horizontal lines of the squares show 95% confidence intervals (CI). *p*-values are given for each compound and for the overall effect.

Consistent with our second hypothesis of abiotic stress amelioration (H2), the decline in BVOC emissions with increasing biodiversity could partly be due to a reduced competition for resources (e.g., light and water) because the emission of BVOCs acts as a defence strategy during plant competition ^66^. Besides, mixed plots can alleviate thermal and water stress ^38,67^ through spatial and structural complexity of canopy cover ^67^ and also differences in rooting depth ^67^. In this context, microclimate buffering is important, as at a global scale, an increase of 2–3 °C in the mean global temperature due to global warming has been shown to increase global BVOC emissions by 30–45% ^68^. Decreasing BVOC emissions with increasing biodiversity also supports our third hypothesis of reduced biotic stress (H3), and can be explained by multiple mechanisms: First, a higher percentage of leaf area eaten by herbivores, which may occur in monoculture due to a higher probability of herbivores^44^, can increase BVOC emissions ^69^. Second, increasing plant diversity decreases herbivore per capita effects on plants while simultaneously benefiting predators ^43,44^. Third, heterogeneity in plant defense strategies in highly diverse communities reduces herbivore performance ^70^. Fourth, plant-herbivore interactions also highly depend on the heterogeneity of available nutrients. For instance, herbivore abundance and performance decrease with plant nutrient variability ^47,48^, indicating that diverse stands are expected to have reduced herbivory damage ^71^, resulting in lower BVOC emissions. Altogether, biotic and abiotic patterns may result in lower emission rates of BVOC in mixtures compared to monocultures, stressing the potential importance of both, biotic and abiotic factors on BVOC emissions^23,72^, yet further experiments are urgently needed to properly test these hypotheses.

### BSOA formation across biodiversity

We quantified a total of fifteen BSOA compounds on the investigated plots throughout the 2021 sampling period. These analytes were diaterpenylic acid acetate [DTAA], 3-methyl-1,2,3-butanetricarboxylic acid [MBTCA], norpinonic acid, pinonic acid, terebic acid, terpenylic acid, pinic acid, adipic acid, pimelic acid, azelaic acid, suberic acid, succinic acid, glutaric acid, salicylic acid, and sebacic acid. Although the observed BSOA compounds did not significantly differ between individual tree species and their corresponding mixtures (Figure 5), succinic and sebacic acid were tendentially higher in monoculture compared to mixture plots (Figure 5L,O); by contrast, pinonic and terebic acid were tendentially higher in mixture plots compared to monoculture plots (Figure 5D,E).

**Figure 5.**
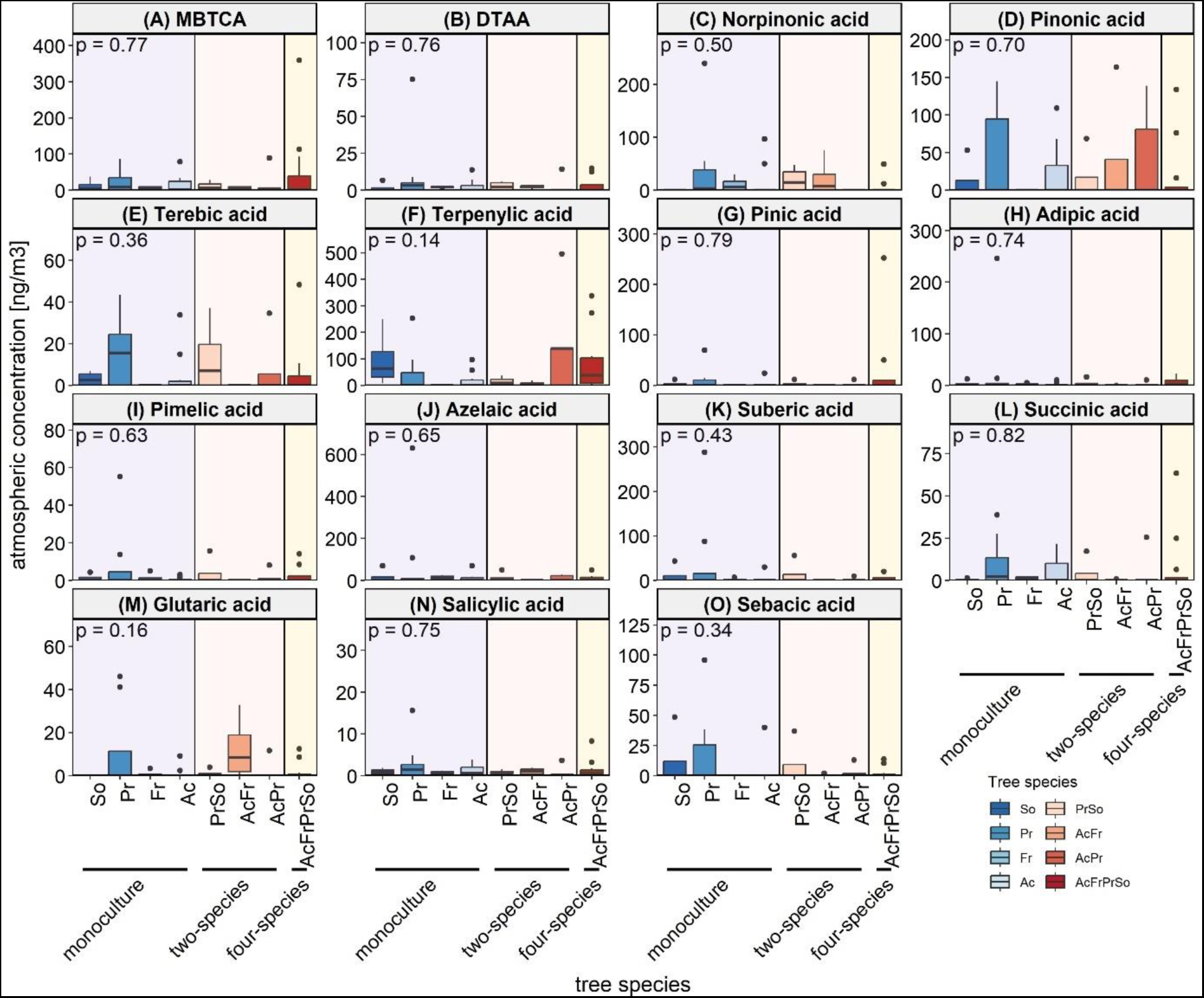
Changes in BSOA compounds across individual tree species and mixtures. Boxes indicate interquartile ranges (bottom and top parts of the box), and median lines (horizontal lines within boxes) and whiskers indicate the minimum and maximum of the measurement. *p*-values are shown. Abbreviations: So, *Sorbus aucuparia* L.; Pr, *Prunus avium* (L.) L.; Ac, *Acer pseudoplatanus* L.; Fr, *Fraxinus excelsior*; DTAA, diaterpenylic acid acetate, and MBTCA, 3-methyl-1,2,3-butanetricarboxylic acid.

While we found no significant differences between observed and expected amounts in BSOA compounds, individual mixtures produced tendentially more or less BSOA compounds than expected. In particular, a mixture of *Acer pseudoplatanus* L. and *Fraxinus excelsior* L. and also a mixture of *Prunus avium* (L.) L. and *Sorbus aucuparia* L. produced a larger than expected amount of norpinonic acid (Figure 6C). Similarly, the mixture of *Acer pseudoplatanus* L. and *Fraxinus excelsior* L. produced a larger than expected amount of glutaric acid (Figure 6M).

**Figure 6.**
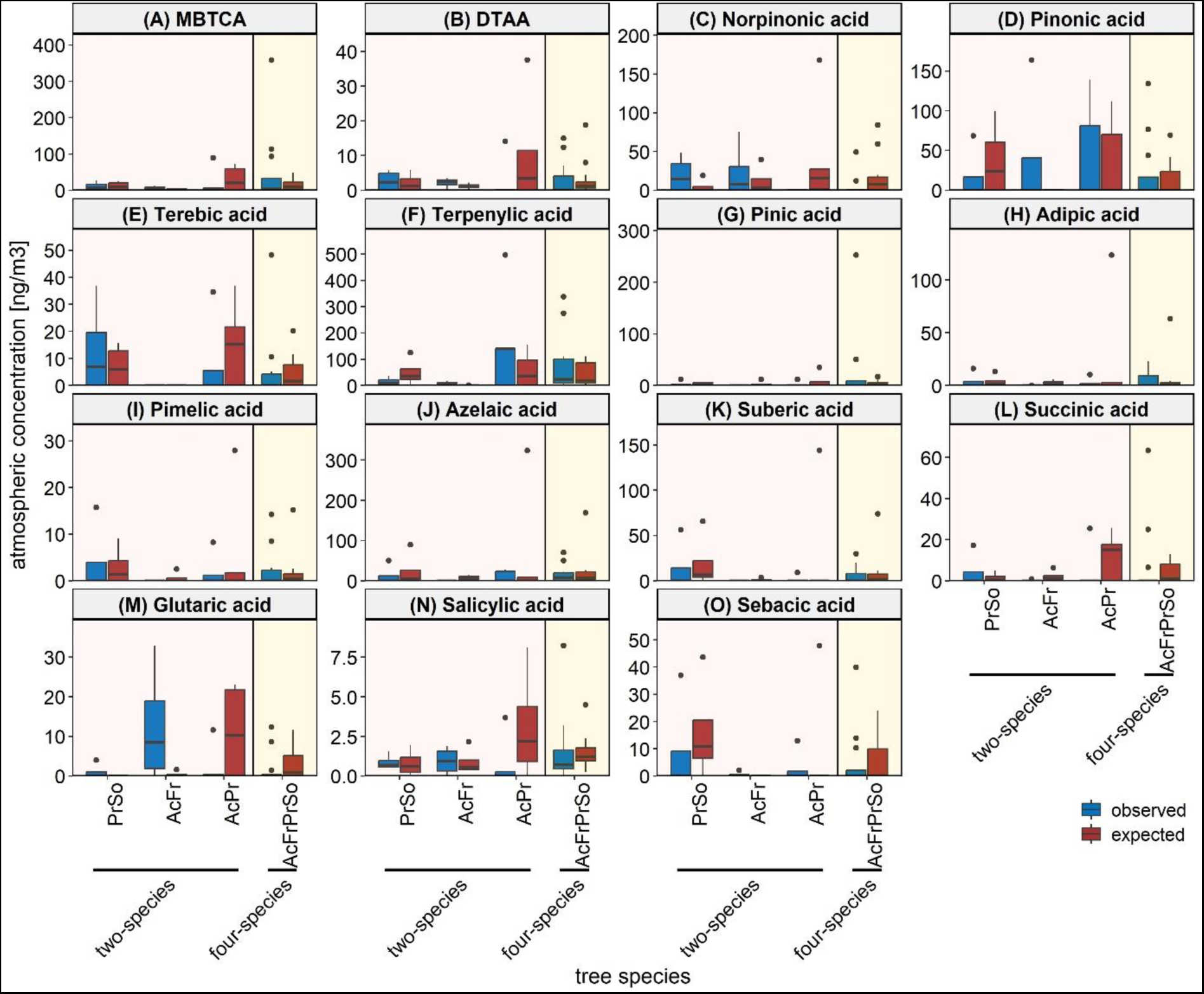
Changes in observed and expected BSOA compounds across tree mixtures (two- and four-species mixtures). Boxes indicate interquartile ranges (bottom and top parts of the box), and median lines (horizontal lines within boxes) and whiskers indicate the minimum and maximum of the measurement. Abbreviations: So, *Sorbus aucuparia* L.; Pr, *Prunus avium* (L.) L.; Ac, *Acer pseudoplatanus* L.; Fr, *Fraxinus excelsior*; DTAA, diaterpenylic acid acetate, and MBTCA, 3-methyl-1,2,3-butanetricarboxylic acid.

Similarly, the relative differences analysis shows that different compounds responded differently to increasing tree diversity, showing that some BSOA compounds (like 3-methyl-1,2,3-butanetricarboxylic acid [MBTCA], pinonic acid, pinic acid, and terpenylic acid) increase relative to what is expected from monocultures but others (like norpinonic acid, adipic acid, suberic acid, and azelaic acid) decrease; however, the overall results were mixed and non-significant (the standardized mean difference: *p* = 0.40, Figure 7). Although the relative differences analysis pattern in two-species and four-species mixtures revealed mixed results, the overall results were significant for two-species mixtures (the standardized mean difference: *p* = 0.05, Supplementary Figure 2A), but non-significant for four-species mixtures (the standardized mean difference: *p* = 0.71, Supplementary Figure 2B).

**Figure 7.**
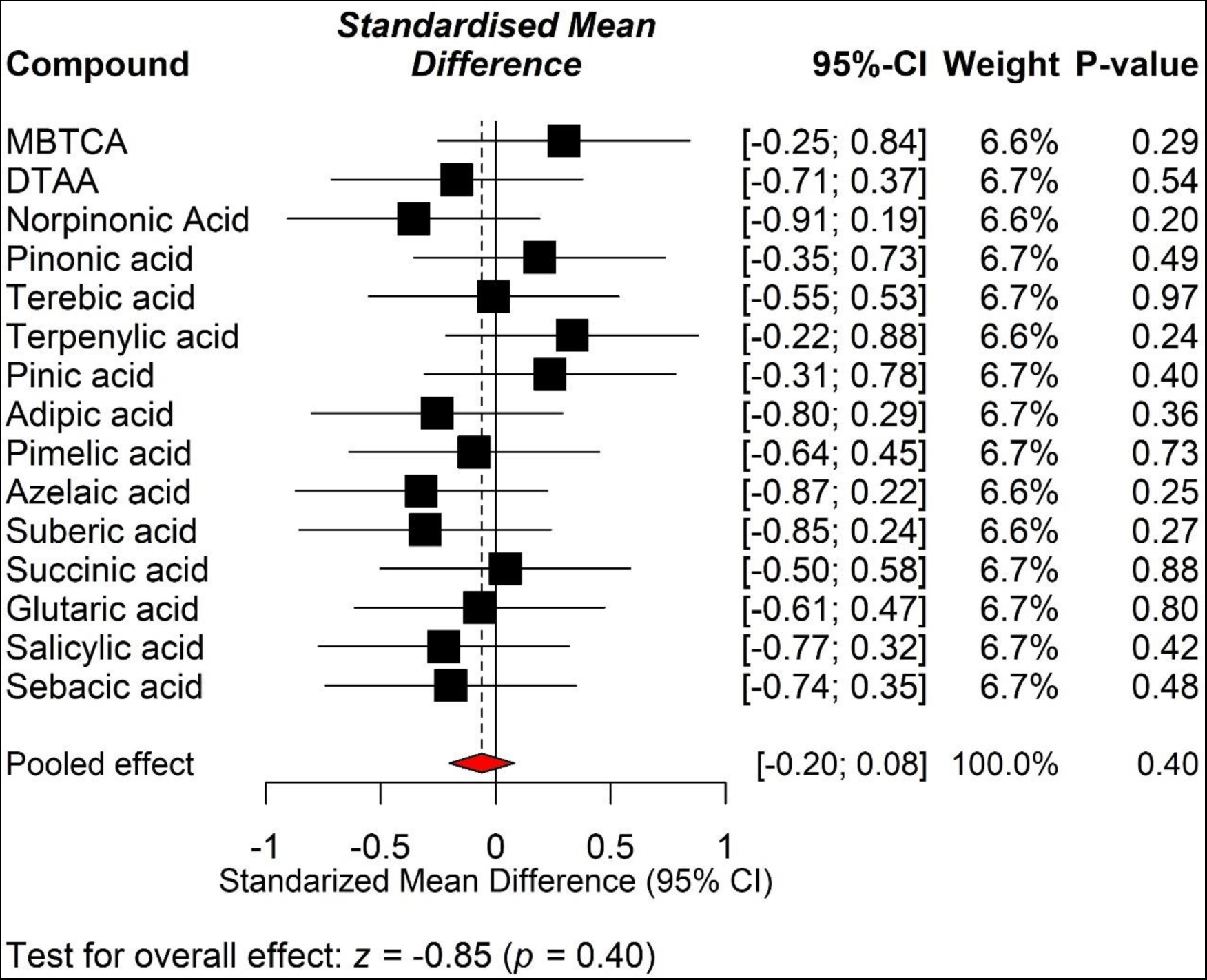
A forest plot summarizing the results of the standardized mean differences between monocultures and mixtures for changes in BSOA compounds as a result of diversity change. The squares’ sizes and widths represent each compound’s weight, while the diamond (red colour) represents the overall effect estimate of the analysis. Effect sizes were calculated using the standardized mean diffrenece. The horizontal lines of the squares show 95% confidence intervals (CI). *p*-values are given for each compound and for the overall effect. Abbreviations: DTAA, Diaterpenylic acid acetate, and MBTCA, 3-methyl-1,2,3-butanetricarboxylic acid.

The mixed results might be due to the influence of regional air masses, as the atmospheric conversions from BVOC to BSOA require some time corresponding to spatial spread and thus preventing the identification of a local BSOA occurrence patterns coupled to the measured BVOC. However, in certain cases, we observed indications of local linkage between atmospheric radicals and α-pinene across individual tree species and mixtures (Supplementary Figure 3). Although our knowledge about biodiversity effects on BSOA formation is limited, it seems likely and it can well be expected that biotic and abiotic stresses also matter indirectly for BSOA formation ^60–62^. For instance, using a compact chamber, temperature dependence of BSOA formation from α-pinene has been reported ^73^, which might partly be related to physical or abiotic differentiation in mixtures versus monocultures. Earlier studies observed a higher formation rate of OH radicals in humid conditions ^74^, which may increase the formation of some BSOA compounds even if BVOCs are low. In addition, the complex formation patterns of BSOA reflect the potential that source precursor and atmospheric oxidants can also matter for oxidation products ^57,58^. Specifically, the differential response of BSOA compounds to diversity gradients could partly be explained by the difference in the gas-phase mechanisms related to the oxidation of α- and β-pinene ^75,76^. For example, MBTCA, which is formed from the oxidation (OH radical) of α-pinene, is a stronger functionalized compound than pinic and pinonic acid ^77^. Overall, we acknowledge that testing our hypothesis needs further experimental work to improve our understanding of how the magnitude and composition of BVOC and BSOA will change across tree diversity gradients.

### Conclusions and research perspectives

In light of significant human-induced changes in biodiversity and climate, the link between atmospheric and biological measurements is crucial to improve our understanding of atmosphere-biosphere feedbacks. However, to what extent changes in atmospheric chemistry and climate change are related to biodiversity is largely unknown. Here we show that the general relationship between biodiversity and BVOC emissions works in principle. The aim of this study is not to properly test our hypotheses, but rather to put forward the relationship between biodiversity and BVOC or BSOA. So, with this first and limited dataset any generalization would be premature. Based on our mixed results, we argue, that measuring BVOC emissions and BSOA formation across biodiversity gradients alone is not enough to fully understand their magnitude and composition. A deeper understanding requires in-depth investigations of microclimate conditions, above- and below-ground herbivores and pathogens, soil microbial communities, accurate monitoring of biotic and abiotic stress, and manipulating biotic and abiotic stress across long-term biodiversity experiments. In addition, we need local and regional-scale models, combined with field and chamber measurements, to improve our understanding of biosphere-atmosphere interactions. These experimental platforms along with the expertise exist in each of the separate disciplines. Therefore, a multidisciplinary approach at the biosphere-atmosphere interface would extend our understanding of the reciprocal effects of biodiversity and climate change ^4^ and help to unravel the extent to which emissions of isoprenoids and aerosol formation are related to biodiversity. This will open a new area of research where the fields of biology, climate science, and atmospheric chemistry may interact.

## Methods

We tested the effect of tree diversity on BVOC emission and BSOA formation by varying tree species richness, including monocultures, two- and four-species mixtures at the MyDiv experimental site located in Saxony-Anhalt, Germany ^65^, between September 21 and October 13, 2021 (13 days in total). The MyDiv experiment comprises eighty 11 × 11 m plots that are located 2 m apart. We used ten of these plots in our case study. The monoculture plots consisted of *Acer pseudoplatanus* L., *Fraxinus excelsior* L., *Prunus avium* (L.) L., and *Sorbus aucuparia* L. The two-species mixture plots consisted of three sets: a mixture of *A. pseudoplatanus* and *F. excelsior*; a mixture of *A. pseudoplatanus* and *P. avium*; and a mixture of *P. avium* and *S. aucuparia*. The four-species mixture plot consisted of all four species. Two monoculture plots, one two-species mixture plot, and one four-species mixture plot were sampled per day for at least four consecutive days (except rainy days). BVOC emissions and BSOA compounds were captured simultaneously using samplers installed at the top of the canopy of each plot. The aluminum sampling box was custom-made and equipped with gas phase (for BVOC) and particulate matter (for BSOA) sampling systems. The gas phase sampling system consisted of a 2-cm-diameter tube with a glass-fritted disc to prevent insect intrusion. The tube was connected to Carbotrap 300 and Tenax cartridges (Supelco, Bad Homburg, Germany) and a GilAir Plus pump (Sensidyne, USA) operating at 150 ml/min using flexible PVC tubes. The particulate matter (PM) system consisted of a PM10 PEM impactor (SKC, USA) inlet connected to a Gillian 12 pump (Sysidyne, USA) operating at 10 l/min. We used an off-line method, where the applied adsorbent cartridges for capturing BVOC and quartz filters for capturing BSOA lasted for four hours (10:00 AM to 2:00 PM). After four hours of sampling, the cartridges and filters were transferred to the laboratory for further analysis. After each day of sampling, the filters were eluted with 3 ml of acetonitrile.

Quantification of BVOC compounds trapped in adsorbent cartridges was then conducted by gas chromatography mass spectrometry (GC/MS, Agilent, Waldbronn, Germany) combined with thermodesorption (Perkin Elmer, Rodgau, Germany). To do so, the BVOCs were first desorbed from the cartridges with a heating rate of 7 °C/min and a flow rate of 50 ml/min to a final temperature of 220 °C. Then the BVOCs were cryofocused at −30 °C on a cooling trap. The trap was heated to 220 °C (hold time: 3 min) with 1 ml/min and the BVOCs were transferred to the Zebron ZB-5ms (30m; 0,25mm; 0,25µm) capillary column. The initial temperature of the column was 35 °C (hold time: 5 min) and then increased stepwise to 120 °C (ramp: 5 °C / min) and finally to 320 °C (ramp: 20 °C / min) and hold for 12 min. Finally, the observed BVOCs were identified using known calibration standards. As a result of this, the following compounds were identified: α-pinene, β-pinene, camphene, Δ3-carene, p-cymene, limonene, α-terpinene, isophorone, and acetophenone.

To analyse BSOA constituents every filter was cut into small pieces and transferred to an extraction vial. 1.0 mL of a water acetonitrile mixture (1:1) was used as an extraction solvent. All samples were shaken for 30 min at 1000 min^-1^ and then transferred into vials with a syringe filter (pore size: 0.2 µm, wwPTFE membrane). The analysis of the extracts was done with ultrahigh-performance LC system equipped with a C_18_ column (Acquity HSST3, 1.8 mm, 2.1 × 100 mm, Waters). The eluents were ultrapure water with 0.1% formic acid (eluent A) and acetonitrile with 0.1% formic acid (eluent B). With a flow rate of 0.3 mL min^−1^, the following gradient was used: 5% B at 0 min, 5% B at 1.0 min, 100% B at 16.0 min, 100% B at 18.0 min, 5% B at 18.1 min, 5% B at 21.0 min. The LC effluent was connected to a high resolution Orbitrap mass spectrometer (Q Exactive Plus, Thermo Fisher Scientific) equipped with an ESI source operated in the negative mode (3.5 kV). Each sample was measured three times in full scan mode (m/z 50−750, R = 70k at m/z 200). Finally, the following BSOA compounds were quantified via an external calibration: diaterpenylic acid acetate [DTAA], 3-methyl-1,2,3-butanetricarboxylic acid [MBTCA], norpinonic acid, pinonic acid, terebic acid, terpenylic acid, pinic acid, adipic acid, pimelic acid, azelaic acid, suberic acid, succinic acid, glutaric acid, salicylic acid, and sebacic acid.

We first compared the amounts of observed emitted BVOC and BSOA compounds across biodiversity gradients using a one-way analysis of variance (ANOVA) test, and where the ANOVA test was significant, we used Tukey’s HSD for pairwise multiple comparisons. Due to the highly reactive nature of α- and β-pinene with atmospheric oxidants such as ozone and OH radical ^78,79^, we performed the Pearson’s correlation coefficients between the products (reacted between α- and β-pinene and ozone and ultraviolet radiation) and the observed BSOA compounds. We analysed the correlation coefficients separately for each measurement week to account for weekly changes of abiotic conditions.

Next, we calculated the expected amount of each BVOC or BSOA compound in the mixture from their respective amounts of observed compounds in monoculture based on the species richness of the mixture plot:

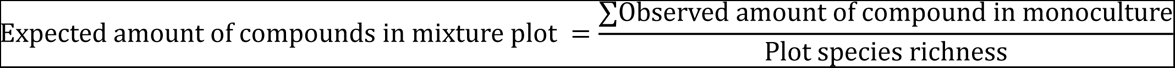

We then compared the observed and expected amounts of each BVOC or BSOA compound using a two-sample t-test.

After that, we used a commonly known Cohen’s d effect size^80^, the standardized mean difference (SMD), to quantify the magnitude and direction of the biodiversity effects on BVOC emission or BSOA formation in monocultures and mixtures. For this, we calculated the SMD for each BVOC compound or BSOA formation and their overall differences between monoculture and mixture using the observed and expected values of each BVOC or BSOA compound ^81^ to see the differences in BVOC emission or BSOA formation between monoculture and mixtures.

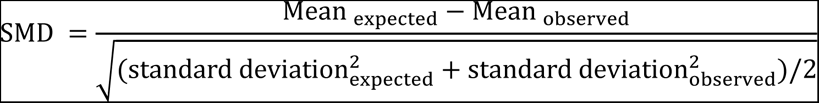

This measure calculates the effect size for each BVOC or BSOA compound, and also for the overall by combining the observed and expected values of all quantified BVOC or BSOA compounds. This approach also assigns a weight to each SMD using the inverse of the variance ^81^ (within-compound variance), which requires the mean, standard deviation as well as the number of observations ^81^ for the observed and expected compounds. To infer how tree diversity levels contribute to BVOC emission or BSOA formation, we performed three separate SMD analyses for BVOC or BSOA compounds for the overall mixtures (a combination of two-and four-species mixtures), two-species mixtures, and four-species mixtures. We finally visualized the results using a forest plot, which shows the effect size of each individual BVOC or BSOA compound and the overall estimates. As such, positive values indicate that the mixture produced a higher amount of compounds than expected based on the monoculture data, and vice versa.

## ACKNOWLEDGMENTS

We thank Sophie Davids and Tom Künne for their help with the field measurements. AS is supported by the Saxon State Ministry for Science, Culture and Tourism (SMWK) – [3-7304/35/6-2021/48880]. HH, TROPOS and TROPOS ACD acknowledge the support of its facilities, some of which have been involved in the present study through ACTRIS-IMP (871115), ATMO-ACCESS (101008004) and ACTRIS-D (01LK2001A). NC was supported by the European Research Council (ERC) under the European Union’s Horizon 2020 research and innovation program (Grant Agreement No. 677232) and by the German Research Foundation (DFG) in the frame of the Gottfried Wilhelm Leibniz Prize (Ei 862/29-1). Further support came from the German Centre for Integrative Biodiversity Research (iDiv) Halle-Jena-Leipzig, funded by the German Research Foundation (FZT 118, 202548816).

## Data availability

We will store the dataset in the public repository once the paper is accepted.

## Competing interests

The authors declare no competing interests.

## Notes

### Competing Interest Statement

The authors have declared no competing interest.

### Summary of Updates

we now have updated the manuscript by adding four more figures in the main file.

