## Supplemental Figures for "Changes in biodiversity impact atmospheric chemistry and climate through plant volatiles and particles"

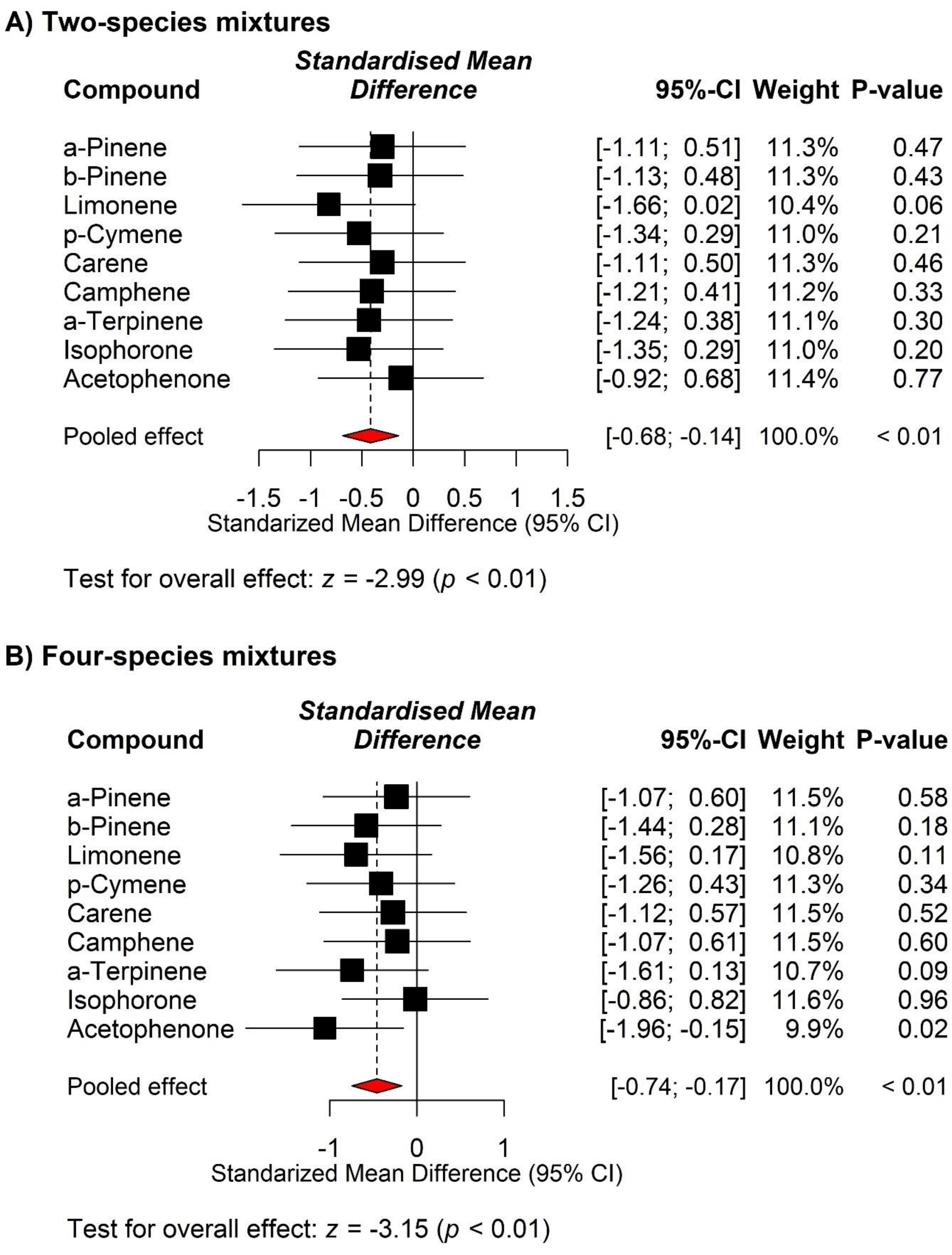


**Supplementary Figure 1.** A forest plot summarizing the results of the standardized mean differences between monocultures and mixtures for changes in biogenic volatile organic compounds (BVOCs) as a result of diversity change in (A) two-species and (B) four-species mixtures. The squares' sizes and widths represent each compound's weight, while the diamond (red colour) represents the overall effect estimate of the analysis. Effect sizes were calculated using standardized mean differences. The horizontal lines of the squares show 95% confidence intervals (CI). *p*-values are given for each compound and for the overall effect.

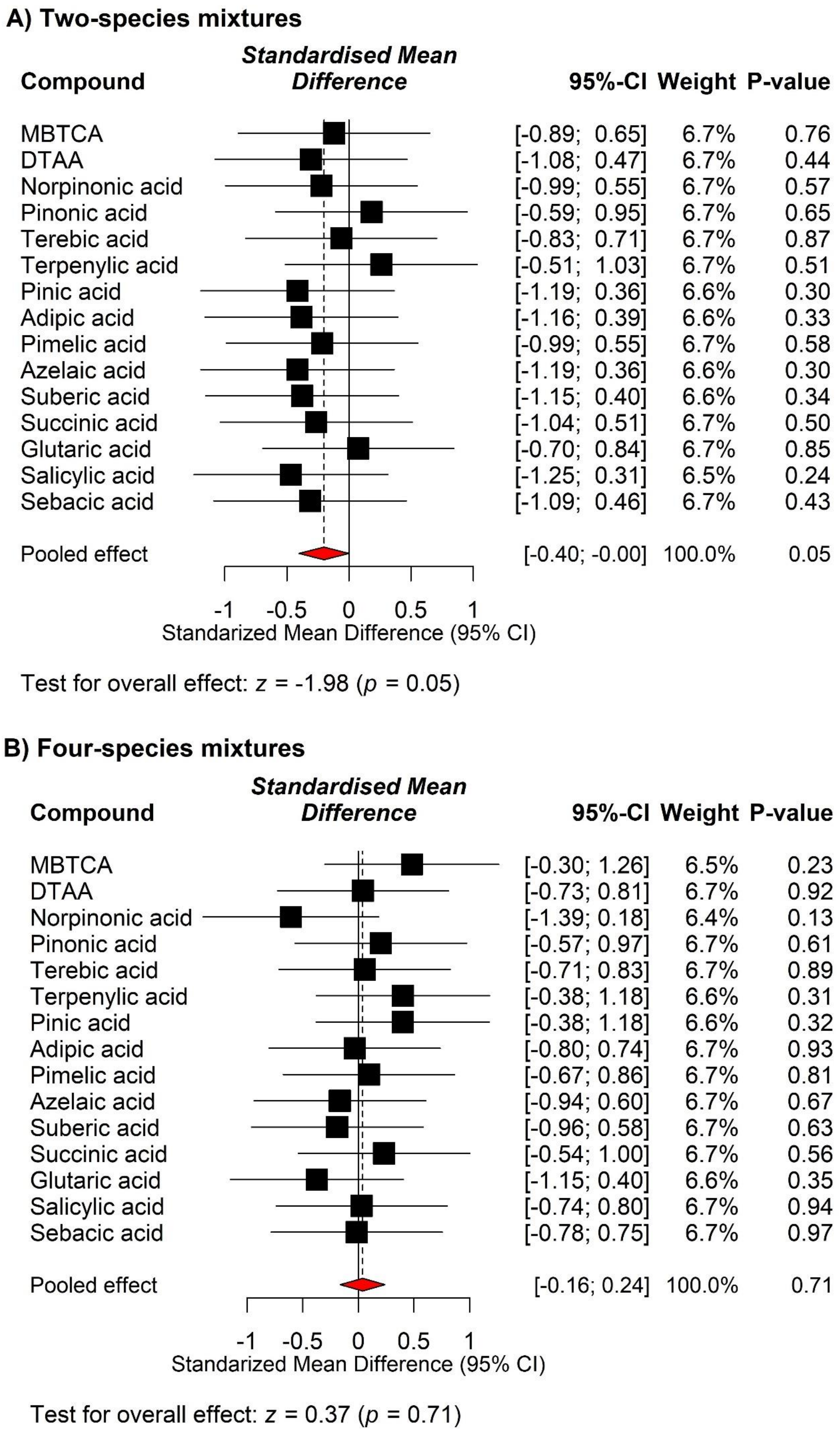
A) Two-species mixtures
Compound
Standardised Mean Difference
95%-CI
Weight
P-value
MBTCA [-0.89; 0.65] 6.7% 0.76
DTAA [-1.08; 0.47] 6.7% 0.44
Norpinonic acid [-0.99; 0.55] 6.7% 0.57
Pinonic acid [-0.59; 0.95] 6.7% 0.65
Terebic acid [-0.83; 0.71] 6.7% 0.87
Terpenylic acid [-0.51; 1.03] 6.7% 0.51
Pinic acid [-1.19; 0.36] 6.6% 0.30
Adipic acid [-1.16; 0.39] 6.6% 0.33
Pimelic acid [-0.99; 0.55] 6.7% 0.58
Azelaic acid [-1.19; 0.36] 6.6% 0.30
Suberic acid [-1.15; 0.40] 6.6% 0.34
Succinic acid [-1.04; 0.51] 6.7% 0.50
Glutaric acid [-0.70; 0.84] 6.7% 0.85
Salicylic acid [-1.25; 0.31] 6.5% 0.24
Sebacic acid [-1.09; 0.46] 6.7% 0.43
Pooled effect [-0.40; -0.00] 100.0% 0.05
-1
-0.5
0
0.5
1
Standarized Mean Difference (95% CI)
Test for overall effect: z = -1.98 (p = 0.05)
B) Four-species mixtures
Compound
Standardised Mean Difference
95%-CI
Weight
P-value
MBTCA [-0.30; 1.26] 6.5% 0.23
DTAA [-0.73; 0.81] 6.7% 0.92
Norpinonic acid [-1.39; 0.18] 6.4% 0.13
Pinonic acid [-0.57; 0.97] 6.7% 0.61
Terebic acid [-0.71; 0.83] 6.7% 0.89
Terpenylic acid [-0.38; 1.18] 6.6% 0.31
Pinic acid [-0.38; 1.18] 6.6% 0.32
Adipic acid [-0.80; 0.74] 6.7% 0.93
Pimelic acid [-0.67; 0.86] 6.7% 0.81
Azelaic acid [-0.94; 0.60] 6.7% 0.67
Suberic acid [-0.96; 0.58] 6.7% 0.63
Succinic acid [-0.54; 1.00] 6.7% 0.56
Glutaric acid [-1.15; 0.40] 6.6% 0.35
Salicylic acid [-0.74; 0.80] 6.7% 0.94
Sebacic acid [-0.78; 0.75] 6.7% 0.97
Pooled effect [-0.16; 0.24] 100.0% 0.71
-1
-0.5
0
0.5
1
Standarized Mean Difference (95% CI)
Test for overall effect: z = 0.37 (p = 0.71)

**Supplementary Figure 2.** A forest plot summarizing the results of the standardized mean differences between monocultures and mixtures for changes in biogenic secondary organic aerosol (BSOA) compounds as a result of diversity change in (A) two-species and (B) four-species mixtures. The squares' sizes and widths represent each compound's weight, while the diamond (red colour) represents the overall effect estimate of the analysis. Effect sizes were calculated using standardized mean differences. The horizontal lines of the squares show 95% confidence intervals (CI). *p*-values are given for each compound and for the overall effect.

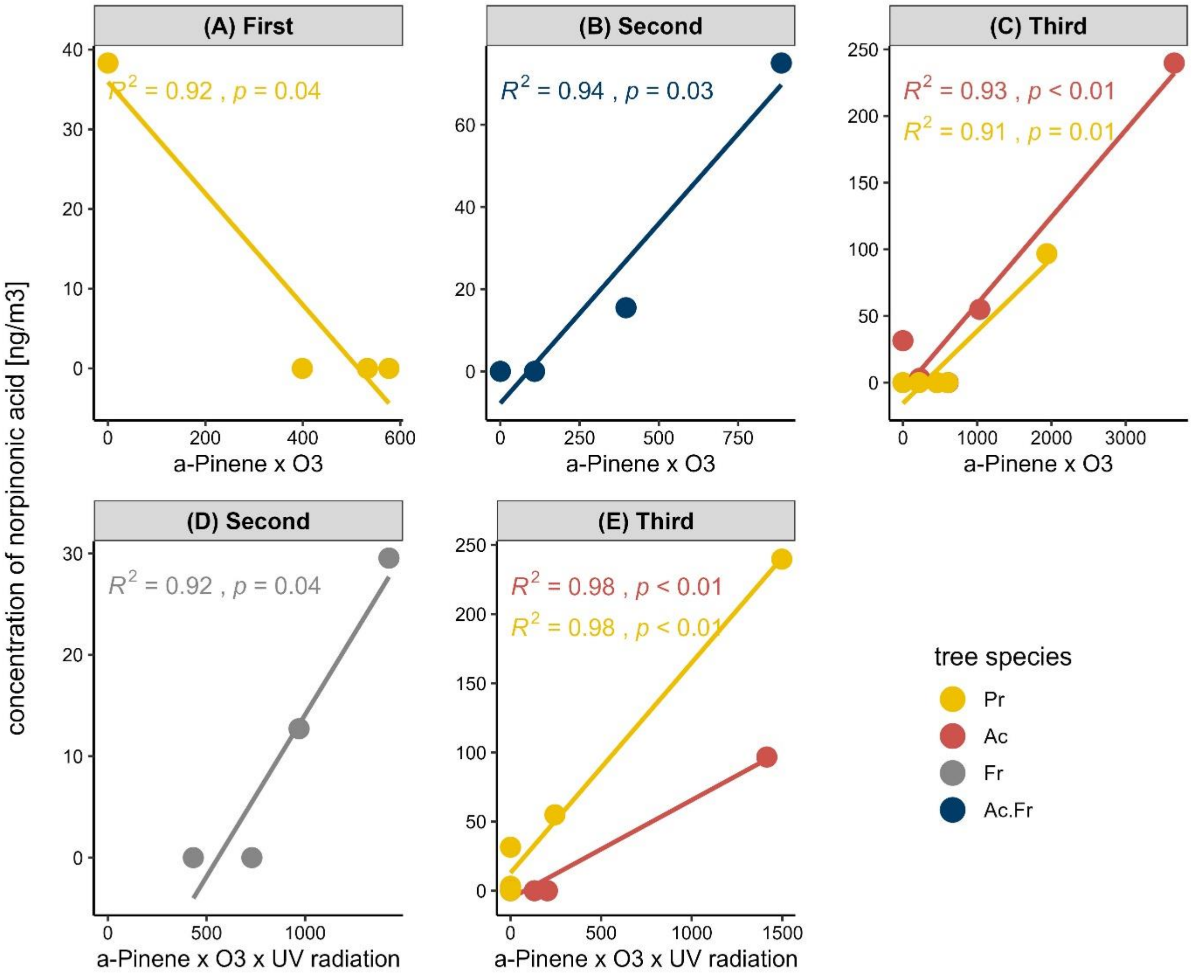


**Supplementary Figure 3.** Correlation between the products and norpinonic acid. A,B,C) α-pinene emission × Ozone D,E) α-pinene emission × Ozone × UV radiation and norpinonic acid across different measurement times (first, second and third weeks) and tree species. *p*-values are shown. Abbreviations: Ac, *Acer pseudoplatanus* L.; Pr, *Prunus avium* (L.) L.; Fr, *Fraxinus excelsior* L., and So, *Sorbus aucuparia* L.
